# Long-read-resolved, ecosystem-wide exploration of nucleotide and structural microdiversity of lake bacterioplankton genomes

**DOI:** 10.1101/2022.03.23.485478

**Authors:** Yusuke Okazaki, Shin-ichi Nakano, Atsushi Toyoda, Hideyuki Tamaki

**Affiliations:** Institute for Chemical Research, Kyoto University, Gokasho, Uji, Kyoto, 611-0011, Japan; Bioproduction Research Institute, National Institute of Advanced Industrial Science and Technology, Central 6, Higashi 1-1-1, Tsukuba, Ibaraki 305-8566, Japan; Center for Ecological Research, Kyoto University, 2-509-3 Hirano, Otsu, Shiga, 520-2113, Japan; Advanced Genomics Center, National Institute of Genetics, 1111 Yata, Mishima City, Shizuoka, 411-8540, Japan

**Author notes:** Corresponding author: Yusuke Okazaki, Institute for Chemical Research, Kyoto University, Gokasho, Uji, Kyoto, 611-0011, Japan.

## Abstract

Reconstruction of metagenome-assembled genomes (MAGs) has become a fundamental approach in microbial ecology. However, an MAG is hardly complete and overlooks genomic microdiversity because metagenomic assembly fails to resolve microvariants among closely related genotypes. Aiming at understanding the universal factors that drive or constrain prokaryotic genome diversification, we performed an ecosystem-wide high-resolution metagenomic exploration of microdiversity by combining spatiotemporal (2 depths × 12 samples) sampling from a pelagic freshwater system, MAG reconstruction using long- and short-read metagenomic sequences, and profiling of single nucleotide variants (SNVs) and structural variants (SVs) through mapping of short and long reads to the MAGs, respectively. We reconstructed 575 MAGs, including 29 circular assemblies, providing high-quality reference genomes of freshwater bacterioplankton. Read mapping against these MAGs identified 100–101,781 SNVs/Mb, 0–305 insertions, 0–467 deletions, 0–41 duplications, and 0–6 inversions for each MAG. Nonsynonymous SNVs were accumulated in genes potentially involved in cell surface structural modification to evade phage recognition. Most (80.2%) deletions overlapped with a gene-coding region, and genes of prokaryotic defense systems were most frequently (>8% of the genes) involved in a deletion. Some such deletions exhibited a monthly shift in their allele frequency, suggesting a rapid turnover of genotypes in response to phage predation. MAGs with extremely low microdiversity were either rare or opportunistic bloomers, suggesting that population persistency is key to their genomic diversification. The results lead to the conclusion that prokaryotic genomic diversification is primarily driven by viral load and constrained by a population bottleneck.

## Introduction

In microbial ecology, reconstruction of metagenome-assembled genomes (MAGs) from an uncultured microbial assemblage has become a routine technique that has reshaped and substantially expanded our understanding of prokaryotic diversity (1, 2). However, MAGs are hardly complete (i.e., circularly assembled) due to difficulties in assembling repetitive (e.g., rRNA genes) and hyper-variable (microdiverse) regions in a genome coexisting in the same sample (3, 4). In particular, genomic microdiversity hampers metagenomic assembly and results in incompleteness or absence of an MAG even at deep sequencing depths, which has been recognized as “the great metagenomics anomaly” (5). Moreover, a metagenomic assembler generally tries to generate a consensus long contig rather than fragmented assemblies reflecting different microvariants (3, 6). Consequently, in a metagenomic assembly, genomic microdiversity is either unassembled or masked by a consensus sequence.

Genomic microdiversity provides information essential to understanding microbial ecology and evolution. Hypervariability of genes involved in cell surface structural modification is thought to be a consequence of the virus–host arms race (7, 8). Intraspecies flexibility of genes responsible for the availability of substrates and nutrients suggests that functionally diversified populations collectively occupy the diverse microniches (9). The degree of genomic microdiversification varies among lineages and is thought to depend on a number of ecological and evolutionary factors such as mutation rate, generation time, population size, genetic mobility, fitness, and drift (10, 11). However, due to the aforementioned difficulties, a comprehensive investigation of genomic microdiversity covering a consortium of microbes in an ecosystem is challenging, and the universal factors that drive or constrain their genomic diversification remain to be elucidated.

To address this, the present study took a three-step approach. The first was comprehensive metagenomic sampling in an ecosystem. We targeted freshwater bacterioplankton assemblages sampled spatiotemporally (2 depths × 12 months) at a pelagic station on Lake Biwa, a monomictic lake with an oxygenated hypolimnion that harbors one of the best-studied freshwater microbial ecosystems (12–16). The second step was long-read metagenomic assembly, which can overcome the problem of fragmented assembly using reads longer than a repeat or hypervariable region (17–20). This was done to generate high-quality reference MAGs covering the diversity of bacterioplankton in the lake. The third step was short- and long-read metagenomic read mapping to the MAGs, in which genomic microvariants were identified as inconsistencies between a consensus assembly and mapped reads (21–23). Notably, we aimed to detect two different types of microvariants, single nucleotide variants (SNVs) and structural variants (SVs), namely, insertion, deletion, duplication, or inversion of a genomic sequence. While short-read mapping efficiently detects SNVs due to its high base accuracy (24, 25), it cannot resolve most SVs that are longer than the canonical short read length (i.e., 150–250 bp). SVs are often associated with gains and losses of genes, which account for a large part of genomic and functional heterogeneity among closely related genotypes (9, 10). Here, the limitation of short-read mapping is complemented by long-read mapping, in which SVs can be located with reads discontinuously aligned to a consensus assembly (26–28). Our three-step approach allowed a high-resolution, ecosystem-wide exploration of SNVs and SVs covering the broad spectrum of prokaryotic diversity in the lake. The results were comparatively analyzed from spatiotemporal, phylogenetic, and gene functionality perspectives, aiming at characterizing factors behind the genomic microdiversification.

## Materials and Methods

### Sample collection

Water samples were collected monthly from May 2018 to April 2019 at a pelagic station (water depth ca. 73 m) on Lake Biwa (35°13’09.5” N, 135°59’44.7” E) from two water depths, representing the epilimnion (5 m) and hypolimnion (65 m) (24 samples in total). Vertical profiles of chlorophyll-a concentration, temperature, and dissolved oxygen were collected using a RINKO CTD profiler (ASTD102; JFE Advantech). The collected lake water was immediately sequentially filtered through a 200 μm mesh, 5 μm polycarbonate filter (TMTP14250; Merck Millipore), and 0.22 μm pore Sterivex cartridge (SVGP01050; Merck Millipore), using a peristaltic pump system onboard. Filtration was performed until the Sterivex cartridge was clogged (1–2.5 liters of lake water were filtered for each cartridge), and at least four Sterivex cartridges were collected for each sample. The filters were flash-frozen in a dry-ice ethanol bath, transported to the laboratory on dry ice, and stored at −80°C until further processing. Water samples were collected between 8:00 am and 11:00 am on each sampling day and processed to the freezing step within 1 h after collection. Prokaryotic cell abundance was determined for each sample using a flow cytometer (CytoFLEX; Beckman Coulter) following fixation of the water sample with 1% glutaraldehyde and staining with 0.25× SYBR Green solution (S7563; Invitrogen).

### DNA extraction

DNA was extracted from the Sterivex filters (i.e., 0.22–5 μm size fraction) using an AllPrep DNA/RNA Mini Kit (80204; Qiagen) with a modified protocol: the filter paper removed from a Sterivex cartridge was put into a Lysing Matrix E tube (6914050; MP Biomedicals) with a mixture of 400 μL RLT plus buffer (containing 1% β-mercaptoethanol following the kit’s protocol) and 400 μL phenol/chloroform/isoamyl alcohol (25:24:1 v/v/v); bead-beating was performed at 3500 rpm for 30 s (MS-100; TOMY Digital Biology), followed by cooling on ice for 1 min, then again at 3500 rpm for 30 sec; the supernatant after centrifugation (16,000 g for 5 min at room temperature) was mixed with 500 μL chloroform/isoamyl alcohol (24:1 v/v) to remove the residual phenol, then centrifuged again; then the supernatant was used as the loading material for the AllPrep DNA spin column and processed following the manufacturer’s instruction. The quantity and quality of the DNA were measured using a Qubit dsDNA HS Assay kit (Q32851; Thermo Fisher Scientific) and a spectrophotometer (NanoDrop 2000; Thermo Fisher Scientific). Consequently, at least 2 μg purified DNA were obtained from each sample.

### Sequencing

The extracted DNA was used for both short- and long-read shotgun metagenomic sequencing. For short-read sequencing, the DNA was sheared to 500 bp on average using an ultrasonicator (Covaris), and a 24-sample multiplexed library was prepared using a MGIEasy Universal DNA Library Prep Set (1000006986; MGI), Circularization Kit (1000005259; MGI), and MGISEQ 2000RS High-throughput Sequencing Set (1000013857; MGI) with seven cycles of PCR amplification. A 1 × 400 bp single-end sequencing was run using one lane of the MGI DNBSEQ-G400 platform. For long-read sequencing, long DNA molecules were purified using diluted (0.45×) AMPure XP beads, and a sequencing library was prepared using a Ligation Sequencing Kit (LSK-109; Oxford Nanopore). Each of the 24 samples was sequenced by an R9.4.1 flow-cell (FLO-MIN106D; Oxford Nanopore) using the Oxford Nanopore GridION platform for 72 h. Base-calling was performed using Guppy (v3.2.10; high accuracy mode).

### Read assembly and contig polishing

Each of the 24 raw long-read libraries was assembled using two different assemblers: Flye (v2.8; – plasmids --meta) (29) and Raven (v1.5.0) (30). The assembled contigs were polished with long reads using Racon (v1.4.13) (31) and Medaka (v1.0.3) (https://github.com/nanoporetech/medaka), and then with short reads using Pilon (v1.23) (32) and two rounds of Racon. Read mapping for polishing was performed using Minimap2 (v2.17) (33) and Bowtie2 (v2.3.5.1) (34). Quality control of short reads was performed using Cutadapt (v2.5) (35) and fastp (v0.20.0) (36). The detailed workflow and parameters are available in Figure S1.

### Binning and bin curation

Contigs longer than 2.5 kb were selected using SeqKit (v0.13.2) (37) and their read coverage across the 24 samples was calculated by mapping the quality-controlled short reads using CoverM (v0.4.0; - m metabat) (https://github.com/wwood/CoverM). The coverage table was input to MetaBAT (v2.12.1) (38) and MaxBin (v2.2.7) (39) to bin the contigs from each of the 24 Flye and Raven assemblies. The resulting 18,621 bins, containing redundancy derived from 24 samples (2 depths × 12 months), two assemblers (Flye and Raven), and two binners (MetaBAT and MaxBin) (Fig. S1), were curated by the following procedures. Bins sharing an average nucleotide identity (ANI) > 95% were clustered using FastANI (v1.31) (40) and the hclust function (method = “average”) of R v4.0.0 (https://www.r-project.org/). This resulted in 3053 bin clusters and 1595 singletons, hereinafter referred to as superbins. Next, bins in the same superbin were merged as follows. First, bin quality score (BQS) was determined as (completeness – [5 × contamination]), referring to the output of checkM (v1.1.3) (41) for each bin. Then, bins derived from the same sample (i.e., only different in the assembler or binner) were merged using quickmerge (v0.3), which bridges gaps in one assembly (acceptor) using sequences of another assembly (donor) based on alignment overlaps (42). Starting from the bin with the highest BQS as an acceptor, bins were iteratively merged by providing a donor bin in the order of BQS. For bins with the same BQS, the bin with fewer contigs was selected in priority. The “--hco” parameter was set to 20, which means that the aligned length should be more than 20 times longer than the unaligned length to merge two contigs. Next, if multiple merged bins in the same superbin (i.e., those from different samples) showed a BQS > 50, they were further merged in the same manner as above. Notably, inter-sample merges did not always generate a better bin than intra-sample merged bins, presumably because of the genomic compositional heterogeneity between samples. Finally, a representative bin was determined for each of the 4648 superbins by selecting the one with the highest BQS among the original and merged bins.

Among the 4648 representative bins, 331 consisted of a single contig. Because quickmerge does not consider genome circularity, we attempted their circularization in the following procedure. First, using nucmer (v3.1) (43), the first and last 50 kb of the contig were aligned against the set of contigs in the same superbin to find a “bridging contig” that may close the gap between the ends. Next, if a bridging contig was found, it was supplied as “new_assembly.fasta” to the circlator (v1.5.5) merge function with the “--ref_end 50000” parameter (44). If the circularization was successful, the contig was rotated to start from a dnaA gene using the circlator fixstart (--min_id 30) function.

Finally, the 4648 representative bins were quality-filtered at BQS > 50, followed by dereplication using dRep (v3.0.1; -comp 0 -con 100 -sa 0.95 --SkipMash --S_algorithm fastANI) (45). The resulting 575 bins were designated as representative/reference metagenome-assembled genomes (rMAGs).

### Analysis of rMAGs

The 575 rMAGs were taxonomically classified using GTDB-Tk (v1.5.0) with the reference data version r202 (46), and the genes were annotated using prokka (v1.14.6) (47) and eggNOGmapper (v2.1.5) (48). Annotated genes were functionally categorized according to KEGG PATHWAY and KEGG BRITE hierarchies (49) assigned to each gene by eggNOGmapper. For further analysis, we selected the top 25 functional categories that covered 33% of the genes. To evaluate the frequency of indel errors that eluded polishing, we followed the idea of the IDEEL software—interrupted open reading frames (ORFs), which are often introduced by a frameshift, were used as an indicator of indel errors (18). Specifically, amino acid sequences of each rMAG predicted by prodigal (v2.6.3) (50) were aligned against the Uniref90 database (release-2020_06) (51) using DIAMOND blastp (v2.0.6; -k 1 - e 1e-5) (52). Based on the results, the proportion of amino acid sequences in which > 90% of the length was aligned to a Uniref90 sequence was determined for each rMAG and designated as the proportion of ORFs aligned > 90% (POA90) score. Coverage-based abundance relative to the total sequenced DNA in each of the 24 samples was determined as reads per kilobase of genome per million reads sequenced (RPKMS), which was generated by mapping the quality-controlled short reads to the 575 rMAGs using bowtie2 (v2.4.2) (34), followed by counting of mapped and unmapped reads using CoverM (--min-read-percent-identity 92). Habitat preference (epilimnion or hypolimnion) of each rMAG was determined using the metric P_epi_, which was defined as the quotient of RPKMS in the epilimnion versus the sum of the value in the epilimnion and hypolimnion (i.e., epilimnion /[epilimnion + hypolimnion]) during the stratification period (May to December). When P_epi_ was > 0.95 or < 0.05, the rMAG was determined as an epilimnion or hypolimnion specialist, respectively (13).

### Analysis of SNVs and SVs

The gene loci and mapping results (i.e., bam files) generated above were input to inStrain (v1.0.0; profile --database_mode --pairing_filter all_reads), which provides genome- and gene-wide SNV profiles based on the short-read alignment (24). SVs were detected by mapping the raw long reads to the rMAGs using NGMLR (v0.2.7) (26) and inputting the resulting bam files to Sniffles (v1.0.12) (26). Among the five types of SVs reported by Sniffles, deletion, insertion, duplication, and inversion were further analyzed, while translocation was removed in the downstream analyses because translocation can involve multiple contigs in different bins and is hard to interpret in metagenomic data. Subsequently, SVs with low (< 0.1) allele frequency (reported by Sniffle) were filtered out. SVs longer than 100 kb were also removed as they were seemingly artifacts introduced by genome circularity, which Sniffles does not account for.

The representative sample providing the highest short-read coverage among the 24 samples was determined for each rMAG, and the result from the representative sample was used for representative SNV and SV profiles. To remove low-quality data derived from low read coverage, rMAGs that showed > 10× short-read coverage in the representative sample (n = 178) were selected and analyzed in detail.

## Results

### General characteristics of the rMAGs

The 24 samples were associated with broad physicochemical conditions. Thermal stratification occurred from May to December, and the prokaryotic cell abundance was 0.82–4.30 (average = 2.00) ×10^6^ cells mL^−1^ (Table S1). For each of the samples, 10.9–27.5-Gb long reads (N50 = 4360–5807 bp) were assembled, and the resulting contigs were polished using 7.0–9.3-Gb short reads (Table S1 and Fig. S1). From the 24 polished contig sets, our pipeline generated 575 nonredundant rMAGs covering 21 phyla of bacteria and archaea (Table S2). The number of contigs, POA90 (indel correction score, see Materials and Methods for detail), and completeness of the rRNA genes all showed better results in rMAGs with higher short-read coverage (Fig. 1a–c). For each of the 24 samples, 45.4–72.1% (mean = 60.4%) of the short-reads were mapped to any of the 575 rMAGs (Fig. S2), indicating that the rMAGs accounted for the majority of the extracted DNA. A ubiquity–abundance plot (Fig. 1d) demonstrated that the rMAGs included common freshwater bacterioplankton lineages known to dominate in Lake Biwa (12, 13, 53). Relative abundance of the rMAGs revealed their diverse distribution pattern across the months and depths (Fig. S3).

**Figure 1.**
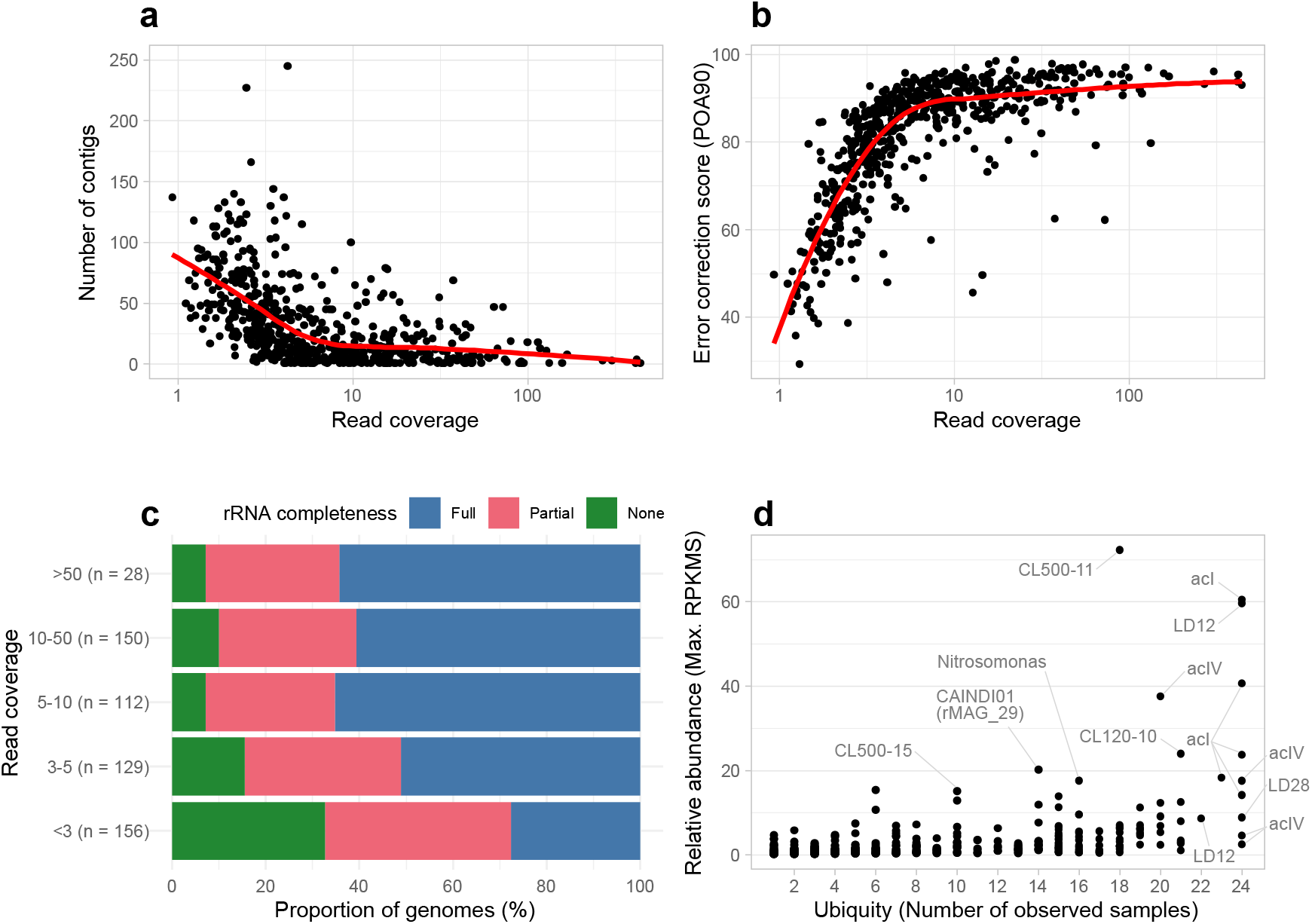
Overview of the 575 rMAGs. Individual rMAGs are represented by each point. Distribution of the (a) number of contigs and (b) error correction score (POA90; proportion of open reading frames [ORFs] aligned > 90% of its length to the reference database) plotted against the read coverage. Solid red lines represent local regression (loess). Read coverage was defined as the average short-read coverage in the representative sample for each rMAG. (c) Proportion of rMAGs with different rRNA gene (i.e., 5S, 16S, and 23S) completeness grouped by read coverage value. (d) Ubiquity–abundance plot of the rMAGs. Relative abundance was defined as maximum reads per kilobase of genome per million reads sequenced (RPKMS) recorded among the 24 samples (i.e., those recorded in the representative sample of the rMAG). Ubiquity was defined as the number of samples in which short reads were mapped to > 50% of the length of the rMAG sequence. Abundant and ubiquitous members are labeled. Detailed statistics for the rMAGs are available in Table S2.

### SNVs and SVs detected in the rMAGs

The 178 rMAGs with > 10× short-read coverage in at least one sample were further analyzed for detection of SNVs and SVs. The results revealed the broad spectrum of genomic microdiversity across the rMAGs (Fig. 2). The number of SNVs per 1 Mb ranged from 100 to 101,781 and significantly varied among the habitat preferences (Fig. 2b). Among the four types of SVs detected, insertion (0– 305 sites per rMAG) and deletion (0–467) dominated over duplication (0–41) and inversion (0–6) (Fig. 2d). The numbers of insertions and deletions were strongly correlated (Pearson’s r = 0.925), while they showed weaker correlations (Pearson’s r = 0.241 and 0.285) with the number of SNVs (Fig. S4). Unlike SNVs, the number of SVs (deletions) did not significantly vary among the habitat preferences (Fig. 2e). Both the numbers of SNVs and SVs (deletions) varied among and within the phyla (Fig. 2c and f).

**Figure 2.**
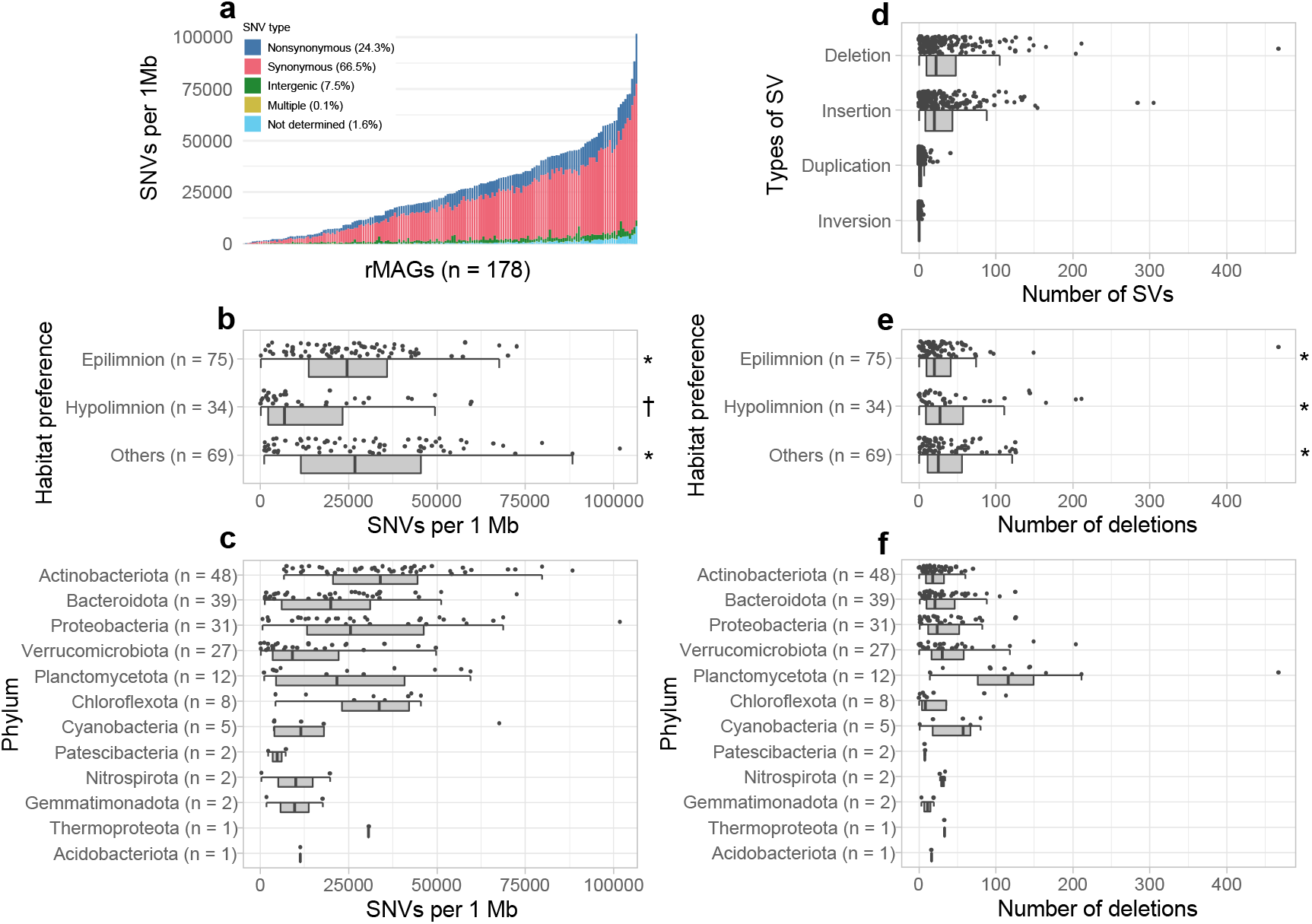
Overview of SNVs and SVs among the 178 rMAGs with > 10× short-read coverage. (a) Each bar represents an individual rMAG, sorted by the number of SNVs per 1 Mb. SNV types determined by inStrain are shown in different colors. The mean proportion of each SNV type among the rMAGs is shown in the color legend. (b–f) Individual rMAGs are represented by each point. Distribution of the number of SNVs per 1 Mb grouped by (b) habitat preference and (c) phylum. (d) Distribution of the number of the four types of SVs in an rMAG. Distribution of the number of deletions in an rMAG grouped by (e) habitat preference and (f) phylum. The same symbol (*or †) in (b) and (e) indicates no significant difference (p > 0.05 in the Wilcoxon rank-sum test) among the groups.

### Genes involved in SNVs and SVs

On average, 66.5%, 24.3%, and 7.5% of SNVs were synonymous, nonsynonymous, and intergenic, respectively (Fig. 2a). The nonsynonymous SNV ratio exhibited a negative correlation with the SNV numbers, and exceptionally high ratios (> 35%) were observed among rMAGs (n = 15) with low SNV numbers (< 7500 per 1 Mb) (Fig. 3a). The nonsynonymous SNV ratio was positively correlated with genome size (Fig. 3b). Gene-resolved SNV frequency and pN/pS exhibited differences among different functional categories (Fig. 4).

**Figure 3.**
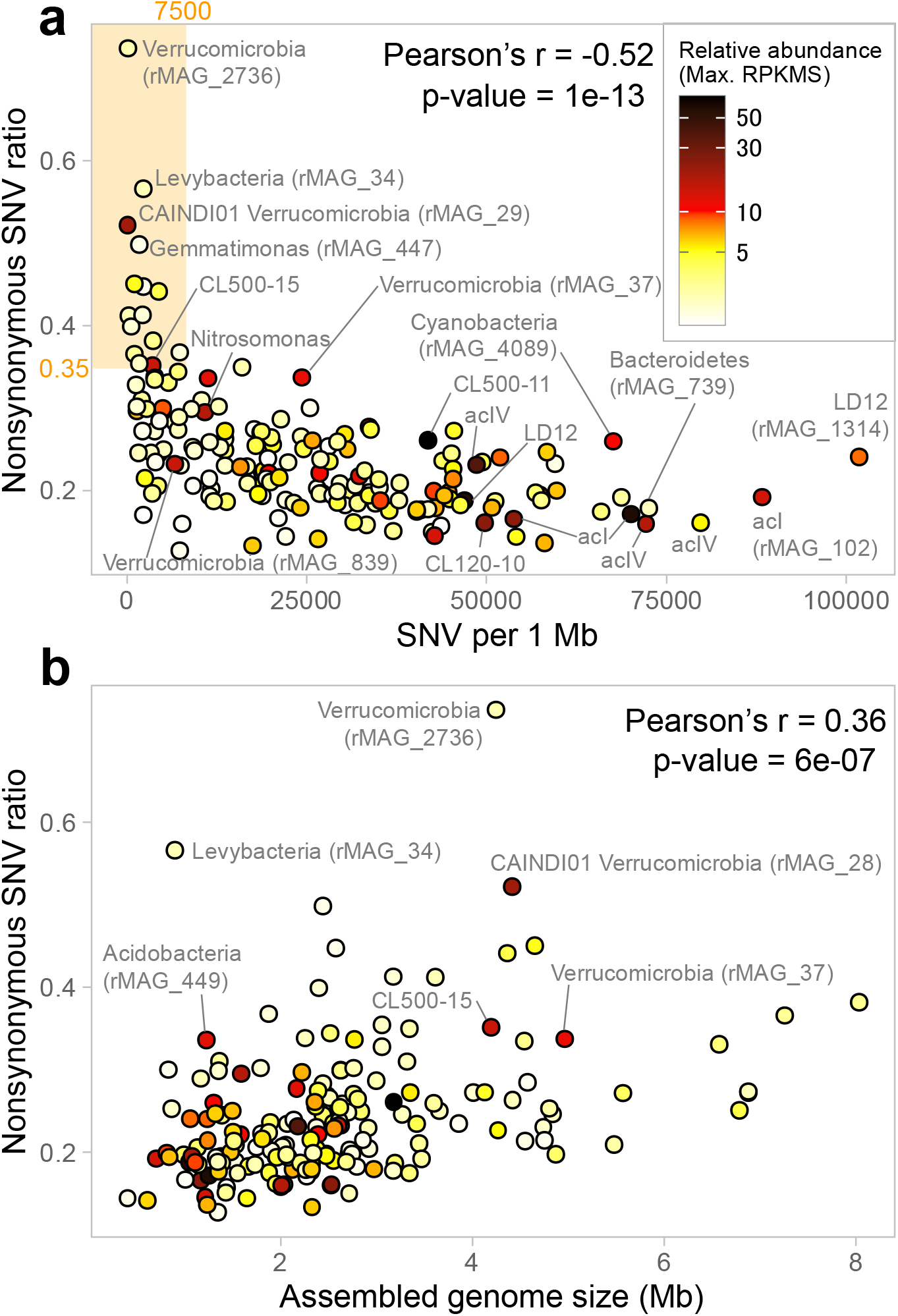
Nonsynonymous SNV ratio of each rMAG plotted against the (a) number of SNV per 1 Mb and (b) assembled genome size. Plot color indicates the relative abundance (maximum RPKMS) of each rMAG defined same as in Figure 1. Representative rMAGs with a high relative abundance or nonsynonymous SNV ratio are labeled. The orange-shaded area on (a) delineates the 15 rMAGs with outstandingly high nonsynonymous SNV ratios (> 35%) and a low number of SNVs (< 7500 per 1 Mb).

**Figure 4.**
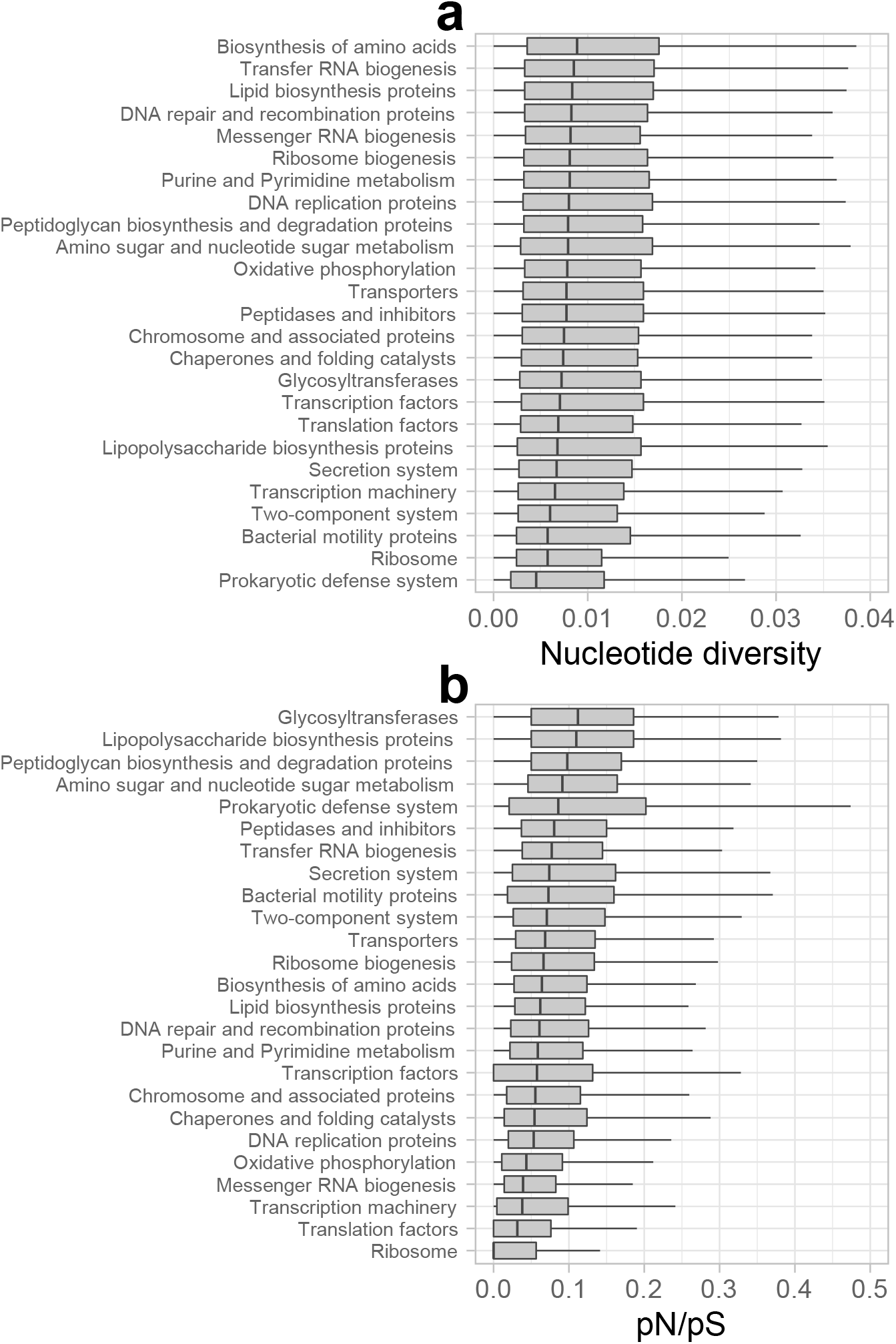
Boxplots indicating the distribution of the (a) nucleotide diversity and (b) pN/pS of genes among the 178 high coverage rMAGs grouped by gene categories. The categories are sorted by the median. Both nucleotide diversity and pN/pS were determined by inStrain. The nucleotide diversity of a gene is defined as a gene-wide average of base-wise nucleotide diversity defined as 1 – (F_A_^2^ + F_C_^2^ + F_G_^2^ + F_T_^2^), where F_X_ is the frequency of base X in the given nucleotide position.

Among the four types of SVs, we further focused on deletions because deletion was the most prevalent SV type (Fig. 2d), and genes involved in deletions can be simply characterized on a genome. The second is not the case for insertion, in which the involved genes appear in the mapped long reads, which are unpolished and unannotated. On average, 80.2% of deletions overlapped with a gene-coding region (Fig. 5a), and the ratio of gene-coding deletions showed a wide range within and among the phyla (Fig. 5b). Gene-coding deletions were most frequently overlapped with transporter genes, which reflects the large number of transporter genes in the rMAGs (Fig. S5). Normalized by the gene counts, genes associated with the prokaryotic defense system were most often (> 8% of the genes) involved in deletions (Fig. 6a). Among the genes affiliated with the prokaryotic defense system, those associated with the type I restriction and modification (RM) system were most abundant in deletion, followed by genes comprising toxin–antitoxin (TA) systems, other RM systems, and CRISPR–Cas systems (Fig. 6b).

**Figure 5.**
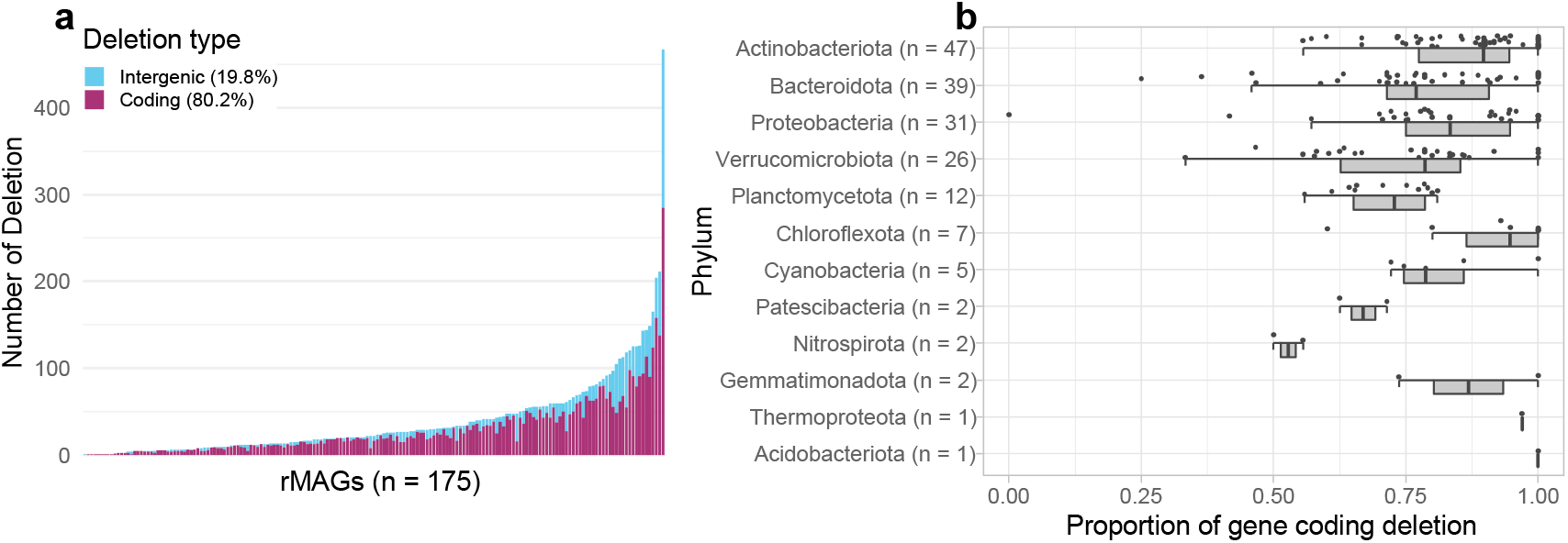
Overview of deletions among rMAGs. Three rMAGs with no deletions were removed from the analysis; the remaining 175 high-coverage rMAGs are shown. (a) Each bar represents an individual rMAG, sorted by the number of deletions. Coding (i.e., overlapping with a gene-coding region) and intergenic deletions are shown in different colors. The mean proportion of each deletion type among the rMAGs is shown in the color legend. (b) Distribution of the proportion of gene-coding deletions grouped by phylum. Individual rMAGs are represented by each point.

**Figure 6.**
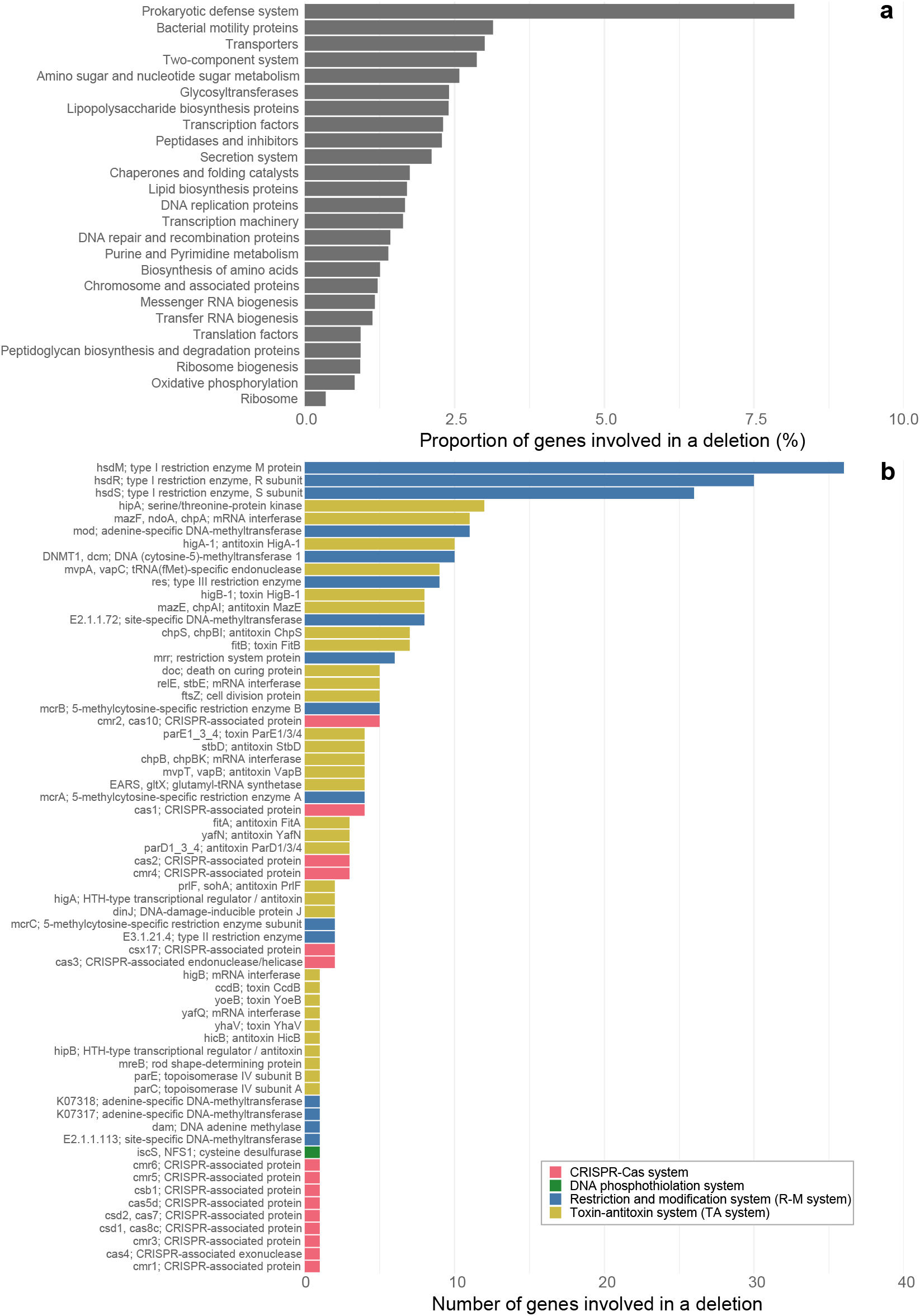
Genes involved in deletions among the 178 high-coverage rMAGs. (a) Proportion of genes involved in a deletion, grouped by gene categories. The same data shown by the number of genes are available in Figure S5. (b) Number of prokaryotic defense system genes involved in a deletion, colored by the type of defense system.

## Discussion

### Long-read metagenomes generated an ecosystem-wide, high-quality prokaryotic genome collection from Lake Biwa

Long-read metagenomics successfully reconstructed high-quality MAGs (Fig. 1) representing the majority of the prokaryotic diversity in the lake across seasons and depths (Fig. 1d and Fig. S2), which was not possible by conventional short-read metagenomics in Lake Biwa (13) or other deep freshwater lakes (54–56). The MAGs included 29 closed assemblies, including the first circular representatives of predominant hypolimnetic bacterioplankton lineages, namely Chloroflexi CL500–11 (rMAG_38), *Nitrosoarchaeum* (rMAG_256), Verrucomicrobia CL120–10 (rMAG_78), Kapabacteria LiUU-9–330 (rMAG_1819), and a member of Nitrosomonadaceae (rMAG_1024) (57, 58).

We should note that we aimed to generate continuous consensus contigs by merging results from different assemblers and samples rather than disjoining microvariants of each genotype. We took this “consensus-first” approach because our subsequent aim was to detect microdiversity masked by the consensus assembly through read mapping. Caveats in analyzing our rMAGs are that they may not represent a single genotype existing in the environment, and they may still contain base errors left unpolished due to inadequate short-read coverage. The POA90 score suggested that fragmented ORFs introduced by uncorrected indel error are common in the majority of genomes with < 10× short-read coverage (Fig. 1b). In light of these limitations, we designate our MAGs as rMAGs (representative/reference MAGs) to differentiate them from those generated by conventional short-read metagenomics and focused on those with > 10× short-read coverage (n = 178) for further analyses.

The general trend that a higher read coverage resulted in a higher-quality rMAG (Fig. 1) suggests that our sequencing effort (Table S1) was unsaturated and deeper sequencing would generate a greater number of high-quality rMAGs. However, read coverage alone was not sufficient to reconstruct a high-quality rMAG. For example, an rMAG of LD12 (*Candidatus* Fonsibacter), which is among the most abundant freshwater bacterioplankton lineages (59, 60), was fragmented and lacked rRNA genes, despite their extremely high read coverage (> 400× in short reads). Members of Pelagibacterales (also known as the SAR11 clade), including LD12, harbor high genomic microdiversity in the flanking region of the rRNA gene operon that is presumably responsible for immunity against their phage (21, 59, 61, 62). Our results indicate that long-read sequencing generally deals well with “the great metagenomics anomaly” (5) but is still unable to solve the issue in extreme cases. Nonetheless, rMAGs provided an unprecedentedly high-quality lake prokaryotic genome collection, which allowed ecosystem-wide exploration of their genomic microdiversity through read mapping.

### Broad spectrum of genomic microdiversity resolved by SNVs and SVs

We found more than 1000-fold differences in the SNV frequency across the rMAGs (Fig. 2a), which is in line with a report on another freshwater system (63). The dominance of synonymous SNVs (Fig. 2a) is also in agreement with previous works in freshwater (63) and marine (21, 64) systems, supporting the idea that the bacterioplankton assemblage is generally under purifying selection with most of the nucleotide variation being neutral. The positive correlation between nonsynonymous SNV ratio and genome size (Fig. 3b) agrees with the hypothesis that genome streamlining is associated with strong purifying selection (65–67). We further found that the frequency of SNVs was lower (Fig. 2b) and also more temporally stable (Fig. S6) in genomes of hypolimnion inhabitants than those of epilimnion inhabitants. These results imply a lower mutation rate in the deeper water layer, presumably due to the lower biological productivity owing to the lower temperature and resource availability in the hypolimnion.

One of the major achievements of the present study was the detection of SVs in a metagenomic sample facilitated by long-read mapping. Compared to the SV analysis for an isolated clonal genome, that for metagenomic assembly generates more complex outputs as it refers to a consensus assembly derived from a highly heterogeneous population. Notably, our approach was not efficient in detecting SVs with a high allele variation or frequency because such a highly heterogeneous region often eludes metagenomic assembly. Given these technical limitations, our goal was not to resolve all SVs, but rather to discover patterns of SV distribution among environmental prokaryotic genomes under the same methodological criteria. Indeed, most SVs in a genome were consecutively detected across samples of different months (Fig. S7), supporting the reproducibility and robustness of our analysis.

Similar to SNVs, we observed significant variation in SV frequency among the rMAGs (Fig. 2d). The relationship between the number of SNVs and SVs was weak because several rMAGs had an extremely high number of SVs (Fig. S4). Notably, members of Planctomycetes harbored disproportionally high numbers of SVs (Fig. 2f) and a lower frequency (55.9–81.0%) of coding deletions (i.e., those overlapping with an ORF) than the average (80.2%) (Fig. 5b). Further investigation found that their non-coding deletions were often associated with intergenic tandem repeats (Fig. S8). Such duplications and deletions can be introduced by slippage of DNA polymerase during replication and can regulate the transcriptional activity or act as a recombination site (68). Planctomycetes generally harbor a large genome with a high number of genes with unknown functions (69). A recent exploration of freshwater Planctomycetes MAGs reported a correlation between their genome size and intergenic nucleotide length (70). Together, their intergenic plasticity might play an essential role in maintaining their genomic integrity. Although characterization of individual SVs is beyond the scope of the present study, overall, our long-read–resolved ecosystem-wide analysis reveals the ubiquity of SVs in environmental prokaryotic genomes and sheds light on their role in regulating genomic structure and function.

### Genetic bottleneck as a major constraint of genomic microdiversity

The negative relationship between SNV frequency and their nonsynonymous rate (Fig. 3a) suggests that stronger purifying selection acts on a genome in which more mutations are accumulated. Based on this assumption, the lineages with a high nonsynonymous SNV ratio and a low number of SNVs may have experienced a recent population bottleneck and not mutated sufficiently to be negatively selected. In other words, their diversification process might still be dominated by random drift or positive selection. Indeed, the top 15 rMAGs with the highest nonsynonymous SNV ratio (delineated in Fig. 3a) were either continuously rare in the hypolimnion or mostly rare but predominant in a short period (boom-and-bust) in either of the water layers (Fig. S3). The former case could be the consequence of the low growth and mutation rates in the hypolimnion, which makes their genome diversification slow enough to be observed before purifying selection dominates. Notably, among these cases, the highest nonsynonymous SNV ratio was observed in rMAG_34, which is affiliated with Levybacteria (OP11), a member of the Candidate phyla radiation (CPR) (71). Recently, a comprehensive exploration of freshwater CPR MAGs (72) reported exceptionally high ANI (99.53%) between Levybacterial MAGs reconstructed from Lake Biwa (13) and Lake Baikal (55) metagenomes. We confirmed that our Levybacterial rMAG also belonged to the same species (ANI > 99.5% to both). Collectively, it is possible that Levybacteria was recently migrated from the Eurasian continent to Lake Biwa, and their genomic microdiversity was still constrained by the genetic bottleneck.

Among the latter (boom-and-bust) cases, prominent examples were two Verrucomicrobial rMAGs (rMAG_2736 and rMAG_29), which had extremely low numbers of SNVs and SVs (Figs. 3a and Table S2) and transiently dominated in the either of the water layers (Fig. S3). Both rMAGs were circular, indicating that long-read metagenomes generate a complete assembly unless hampered by high microdiversity or low read coverage. The boom-and-bust dynamics of Verrucomicrobia agrees with the general assumption that they are opportunistic strategists rapidly responding to a supply of carbohydrates (73, 74). Notably, rMAG_29 (taxonomically assigned to the genus “CAINDI01” by GTDB) was among the most abundant bacterioplankton lineages in the lake during their bloom (Figs. 1d and S3), with their relative abundance (RPKMS) increasing over 12-fold in just 1 month (1.39 in November to 16.92 in December). Because their bloom was observed from May to June and from December to January in the hypolimnion (Fig. S3), their growth was likely triggered by a supply of polysaccharides exudated from sinking phytoplankton cells derived from the spring and autumn algal blooms in the epilimnion, as observed in a previous study in the lake (75). Taken together, the ecological strategy of CAINDI01 (to rapidly exploit intermittent resources) produced periodic genetic bottlenecks and effectively eluded selective processes, which resulted in their extremely low genomic microdiversity in the lake despite their quantitative dominance. Interestingly, CAINDI01 encoded as many as 236 transposase genes (annotated by prokka), but none of them were associated with SVs, except for an inversion involving IS21 transposases (data not shown). The results further suggest that their rapid population turnover prevented invasions of mobile genetic elements (MGEs). Collectively, we conclude that a genetic bottleneck is a primary factor constraining genomic microdiversification.

Conversely, the extent of genomic microdiversification may be used to predict the existence or absence of a recent bottleneck event. For instance, rMAG_739 (Chitinophagaceae of the phylum Bacteroidetes) was the fourth-most SNV-rich rMAG, with a low nonsynonymous rate (Fig. 3a), despite the fact that they were detectable only from June to October in the epilimnion (Fig. S3). These results suggest that they did not experience a recent genetic bottleneck and thus are allochthonous, presumably maintaining their large genetic pool in the inflowing river, sediment, or the water column horizontally distant from our sampling site. It should also be noted that no sign of a recent bottleneck event was found among common and abundant freshwater bacterioplankton lineages (e.g., LD12, acI, acIV, and CL500–11). Interestingly, the two most SNV-rich members, rMAG_1314 and rMAG_102, were continuously and ubiquitously abundant species of LD12 and acI, respectively, rather than the most abundant ones (i.e., rMAG_300 and rMAG_28) of the lineage (Figs. 3a and S3). These facts further support the hypothesis that persistent rather than abundant populations exhibit higher intra-population sequence variation (76).

### Phage predation as a major driving force of genomic microdiversification

The lowest pN/pS in housekeeping genes involved in replication, transcription, translation, and oxidative phosphorylation (Fig. 4b) agreed with a previous study in the Baltic Sea (25) and indicated that genes involved in core functions are under stronger purifying selection. By contrast, high pN/pS were observed among genes potentially involved in cell surface structural modification, namely glycosyltransferases, lipopolysaccharide biosynthesis, and peptidoglycan biosynthesis proteins (Fig. 4b). Hypervariability of such genes has been observed in genomes of ubiquitous marine and freshwater bacterioplankton and is considered beneficial in evading the host recognition system of their phage (7–9). Our results further demonstrate that these traits are universal in the ecosystem and suggest that phage predation is the most prevalent selective pressure generating amino acid-level gene diversity.

The SV profiling demonstrated that deletion was overrepresented in genes involved in prokaryotic defense systems, namely, RM systems, TA systems, and CRISPR–Cas systems (Fig. 6a). Among them, the three proteins making up the Type I RM system (R, M, and S) were the most represented (Fig. 6b). A previous metaepigenomic exploration revealed the diversity of DNA methylated motifs and methyltransferase genes among Lake Biwa bacterioplankton assemblages (77). Interestingly, the study reported a corresponding pair of a methylated motif and a methyltransferase gene is often absent in MAGs, which could be attributable to the incompleteness of MAGs or to the limited sensitivity of the method. Further, the study found that the ratio of methylation in each motif in a genome varied considerably from 19% to 99%, for which the authors reasoned the methodological limitation of modification detection power (77). Our results introduce another possible explanation for these observations: the mobility of RM-related genes within a sequence-discrete population might have resulted in the heterogeneous recovery of methylated motifs or methyltransferase genes in an MAG. The variable nature of epigenetic modification proposes another layer of genomic microdiversity, which will be key to revealing the mechanism behind the virus–host arms race.

The next most represented defense genes in deletions were those involved in TA systems (Fig. 6b), which can also act as an antiphage system (78). Recent experimental work has demonstrated that mobility and rapid turnover of genes involved in intracellular defense machinery are essential mechanisms to maintaining the core genome in the face of phage predation (79). Our results that RM and TA systems are highly mobile (Fig. 6b) suggest the prevalence of such mechanisms in the ecosystem. In addition, SNV analysis revealed that the prokaryotic defense system was the gene category with the lowest nucleotide diversity (Fig. 4a) and among the highest pN/pS ratios (Fig. 4b), which implies that the defense genes are positively selected by phage predation. Meanwhile, both RM and TA systems can behave as selfish and addictive elements and are prone to be horizontally transferred with an MGE (78, 80, 81). Their beneficial and parasitic aspects are not mutually exclusive, and the relative contribution of the two remains unresolved. Thus, we cannot rule out the possibility that some defense genes are rather parasitic and nonbeneficial or even detrimental for the host. In any case, these genes are among the most prevalent mobile genes generating genomic heterogeneity within a sequence-discrete population.

Although not as frequent as RM and TA systems, we also found deletions associated with genes involved in the CRISPR–Cas system (Fig. 6b). Further investigation revealed individual cases in which the whole CRISPR–Cas system was involved in a deletion, and one of them further included TA system genes (Fig. S9). Experimental studies have suggested that the CRISPR–Cas system can disseminate horizontally (82, 83) and is sometimes encoded in an MGE, which facilitates not only adaptive immunity against phages but also inter-MGE competition and guided transposition of the MGE (84–86). Our results provide evidence of the mobility of the CRISPR–Cas system in an ecosystem, although it remains unknown whether it is beneficial or parasitic for the host.

Finally, we note that our monthly investigation revealed a shift in the allele frequency of deletions or insertions involving the CRISPR–Cas system and CRISPR spacers during the study period (Figs. S9 and S10). The results suggest monthly turnover of the population composition driven by the virus–host arms race. Such a rapid shift of population composition has been demonstrated from the virus side in the marine system (22). Our results are the demonstration from the host side and propose the significance of not only sympatric but also temporal microdiversity. In summary, our ecosystem-wide investigation of SNVs and SVs suggests that phage predation is the major driving force of genomic microdiversification among the environmental microbial assemblage. The key question for future works is whether and how the mobility of defense genes is beneficial for the host, for which the microdiversity of the counteracting viral genome must be explored.

## Conclusion

Our ecosystem-wide high-resolution approach combining spatiotemporal sampling and long- and short-read metagenomics resulted in two major achievements. First, we presented a collection of high-quality MAGs covering the majority of the prokaryotic diversity in a deep freshwater lake, which will be a valuable reference for future studies in freshwater microbial ecology. Then the broad spectrum of SNVs and SVs masked in the MAGs were detected by short- and long-read mapping, respectively, which is the second and greater achievement of this work. Based on the results, we conclude that genomic microdiversification is primarily driven by viral load and constrained by genetic bottlenecks.

We also demonstrated the performance and limitation of our “consensus-first” approach (Fig. 1). To push the consensus-first approach further, future works can consider gaining a deeper sequencing depth (for instance, using the PromethION platform (87, 88)) and obtaining longer sequencing reads with a more sophisticated DNA extraction method (89). Alternative possible approaches include genome-free metagenomics, which directly handles pan-metagenomic graphs without the prerequisite of a linear genomic assembly (90). The ultimate approach will be a strain-resolved assembly, which usually requires an isolated culture or single cell but was recently accomplished in a metagenomic assembly using highly accurate long reads (i.e., PacBio HiFi reads) (20), although it is still too costly for common application.

Lakes are physically separated unique ecosystems and thus harbor genetically isolated microbiomes (91), while those in the marine system are likely distributed globally (64, 92) presumably following the rapid circulation of global surface seawater (93). This implies that we can further perform a comparative study among different lakes, in which each lake can be considered as a replicate or control of an ecosystem. The two main factors affecting genome microdiversification (genetic bottlenecks and virus–host interactions) are both lake-specific. The microbiomes in different lakes have a different history of biological interactions in different physicochemical conditions, which would result in different trajectories of genome microdiversification. For instance, we hypothesize that a larger and older lake is less affected by genetic bottlenecks in terms of time and space. That is, the extent of bacterioplankton microdiversification in Lake Biwa (the largest and oldest lake in Japan) might be the greatest among the lakes in the country but might be lower than that of Lake Baikal, the largest and oldest freshwater lake on the earth. Such inter-lake comparative analyses will be an effective approach to further validate the findings in the present study and unveil the universal mechanisms in the diversification and evolution of the microbial genome.

## Data availability

The raw sequencing reads generated in the present study are available under accession numbers DRR333363–DRR333410 (BioProject ID: PRJDB12736) as summarized in Table S1. Nucleotide fasta files of the rMAGs are available in https://doi.org/10.6084/m9.figshare.19165673.v1

## Author contributions

YO and HT conceived the study and performed experimental work. YO and SN performed field sampling. AT performed DNA sequencing. YO conducted data analysis and wrote the draft. All authors contributed to finalizing the draft and approved for the final version.

## Acknowledgements

We are grateful to Yukiko Goda, Tetsuji Akatsuka, Yasuhiko Yamaguchi, and crews of research vessels Hasu, Hakkengo, and Biwakaze for their assistance in Lake Biwa sampling. We thank Bioengineering Lab. Co., Ltd for providing sequencing resources. Computation time was provided by the SuperComputer System, Institute for Chemical Research, Kyoto University. This study was supported by Center for Ecological Research, Kyoto University, a Joint Usage / Research Center, The Kyoto University Foundation, and JSPS KAKENHI Grant Numbers 16H06279 (PAGS), 18J00300, and 19H03302.

