## Supplementary information for "Long-read-resolved, ecosystem-wide exploration of nucleotide and structural microdiversity of lake bacterioplankton genomes"

**Corresponding author:**

Yusuke Okazaki

**This PDF file includes:**

Figures S1 to S10

Captions for Tables S1 and S2

**Other supplementary materials for this manuscript include the following:**

Tables S1 and S2

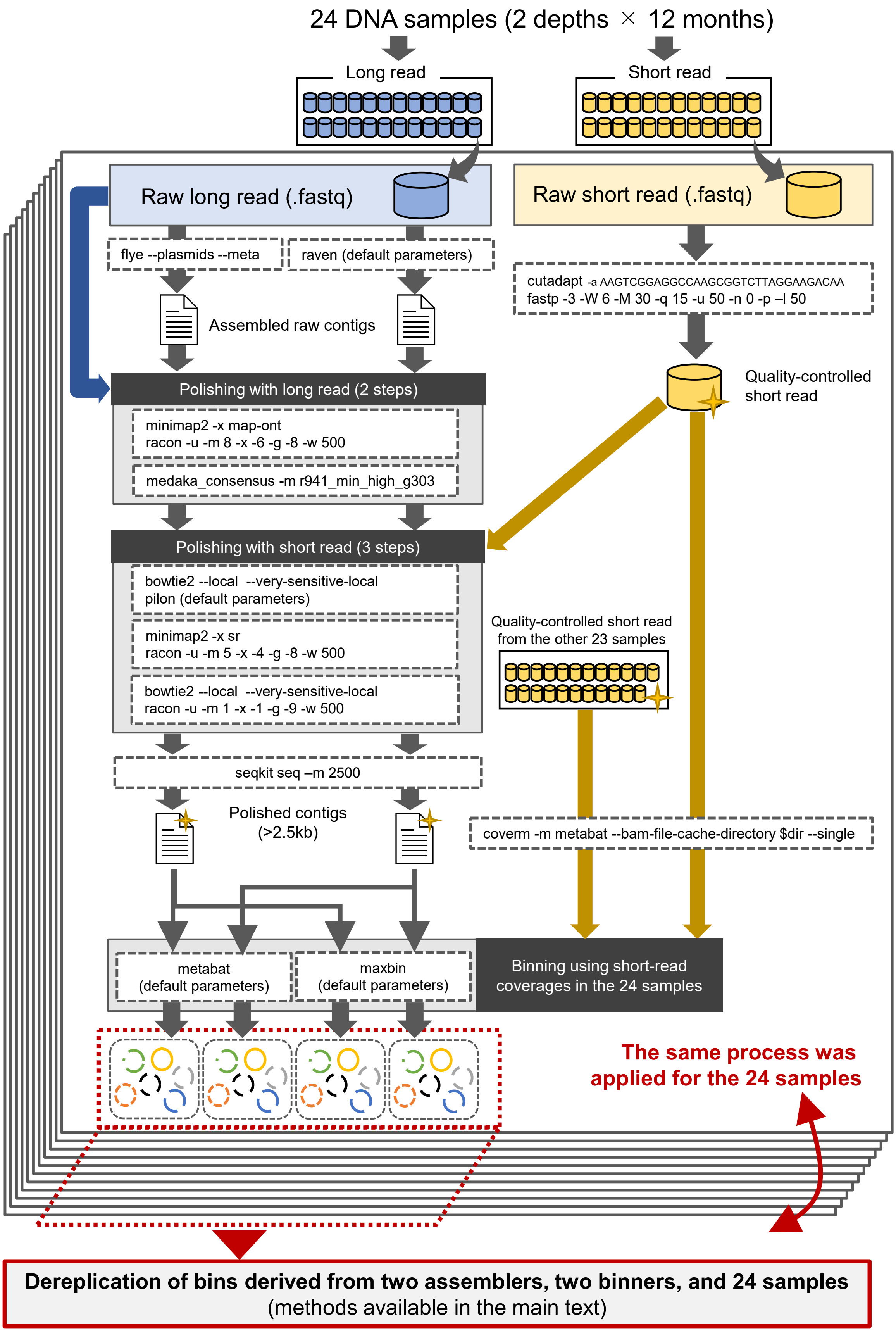


**Figure S1.** Assembly and binning pipeline in the present study. Only parameters required to reproduce the analysis are shown.

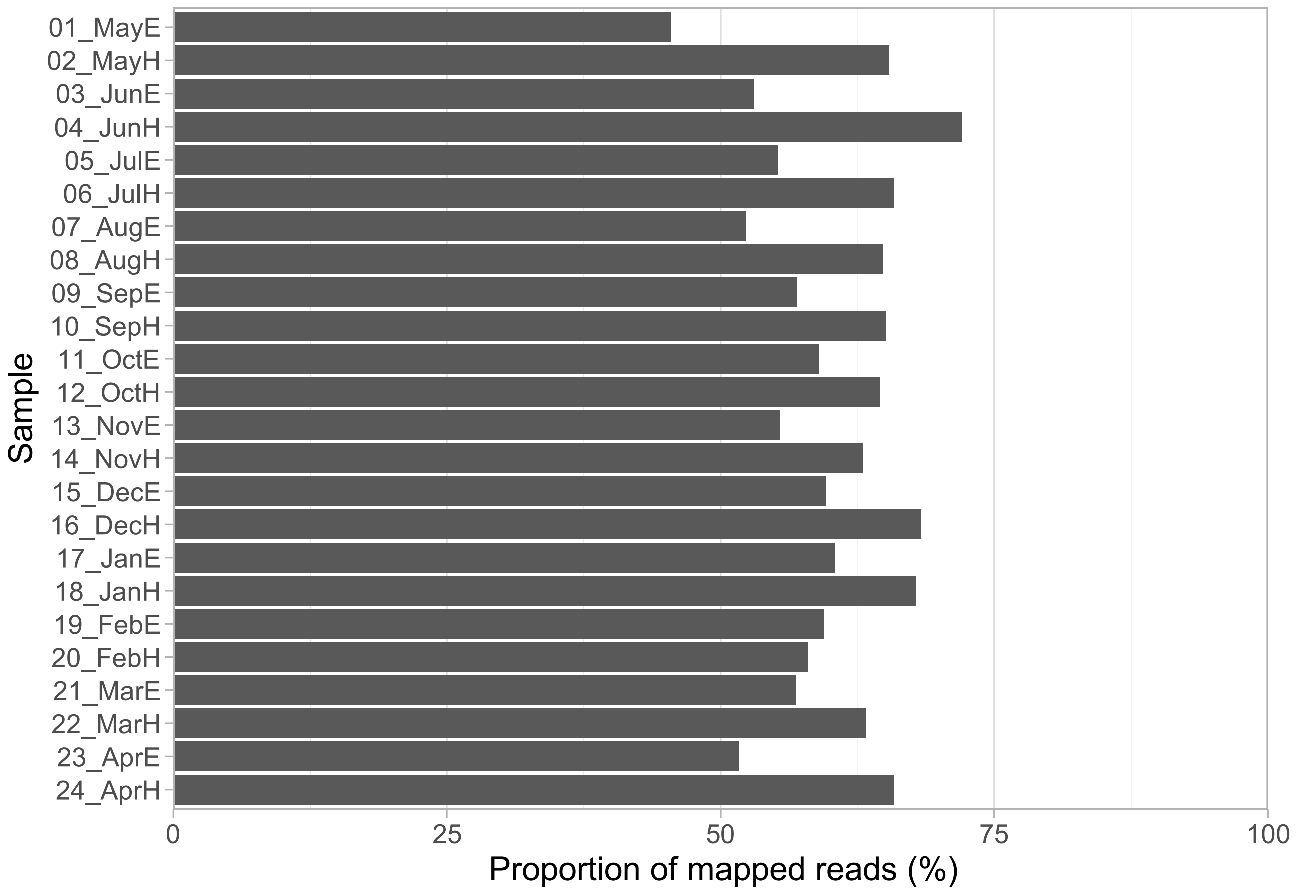


**Figure S2.** Proportion of short reads mapped to the 575 rMAGs.

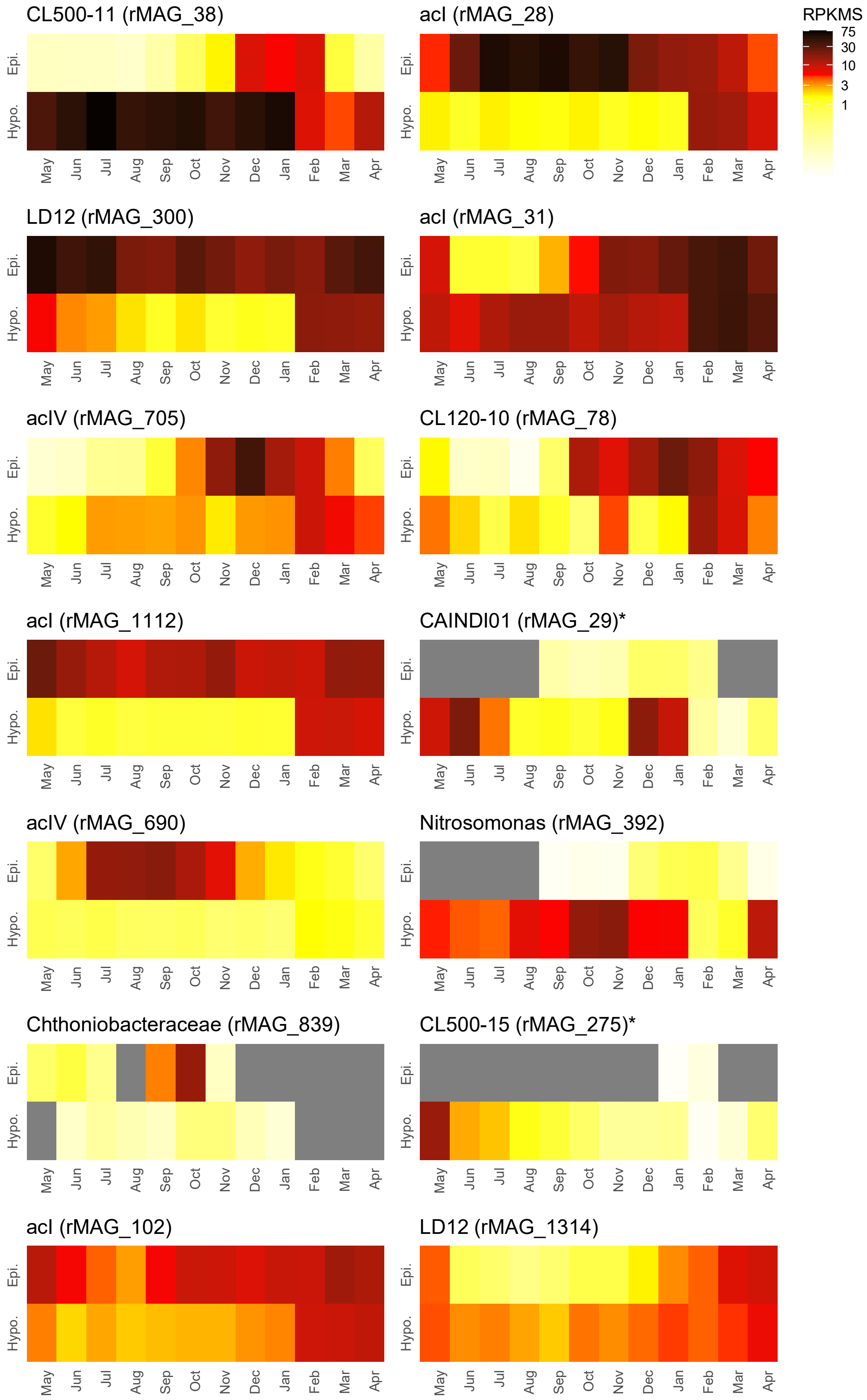


**Figure S3.** Relative abundances (RPKMS) of representative rMAGs across the 24 samples. Asterisks indicate the top 15 rMAGs with the highest nonsynonymous SNV ratios (delineated in Fig. 3a; n = 15). Gray cells indicate RPKMS = 0 (i.e., not detected). The stratification period was from May to December. Epi., Epilimnion; Hypo., Hypolimnion.

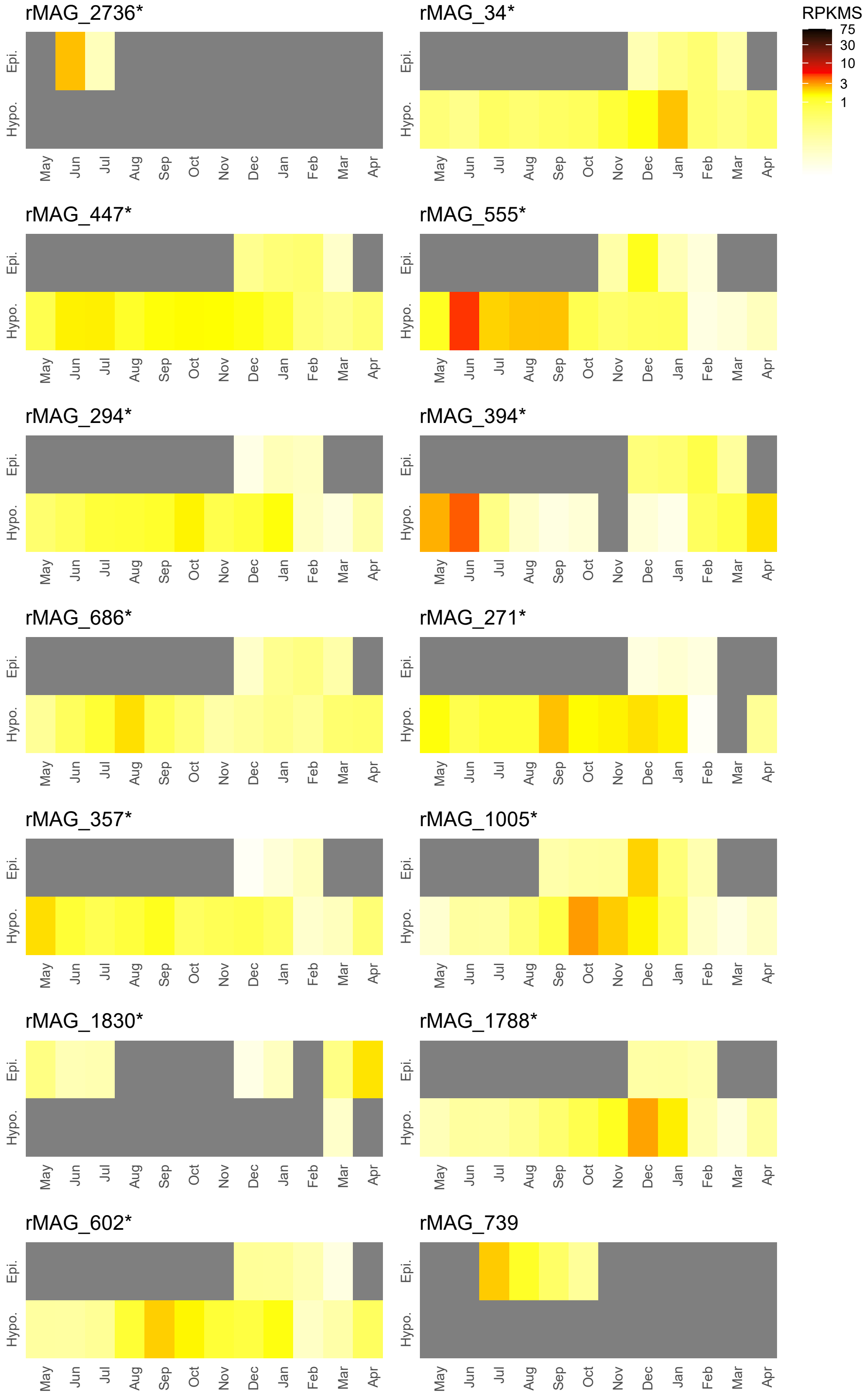


**Figure S3.** (Continued.)

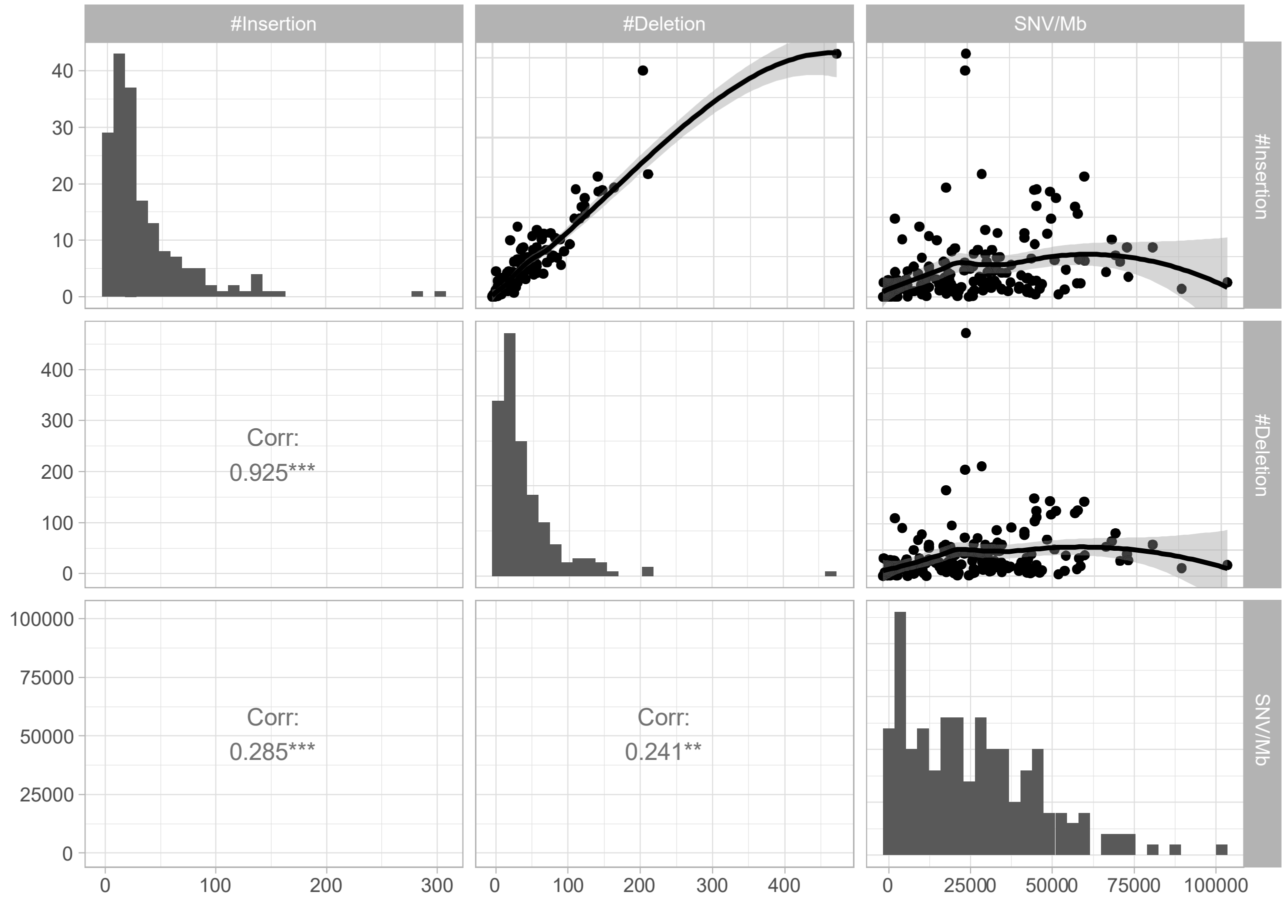


**Figure S4.** Pairwise plots (upper right panels) among the number of insertions, deletions, and SNVs per 1 Mb. Solid line represents local regression (loess); 95% confidence intervals are shaded gray. Histograms on diagonal panels indicate the distribution of each parameter. Bottom left panels show the Pearson correlation (r) with *** and ** indicating p-values of < 0.001 and < 0.01, respectively.

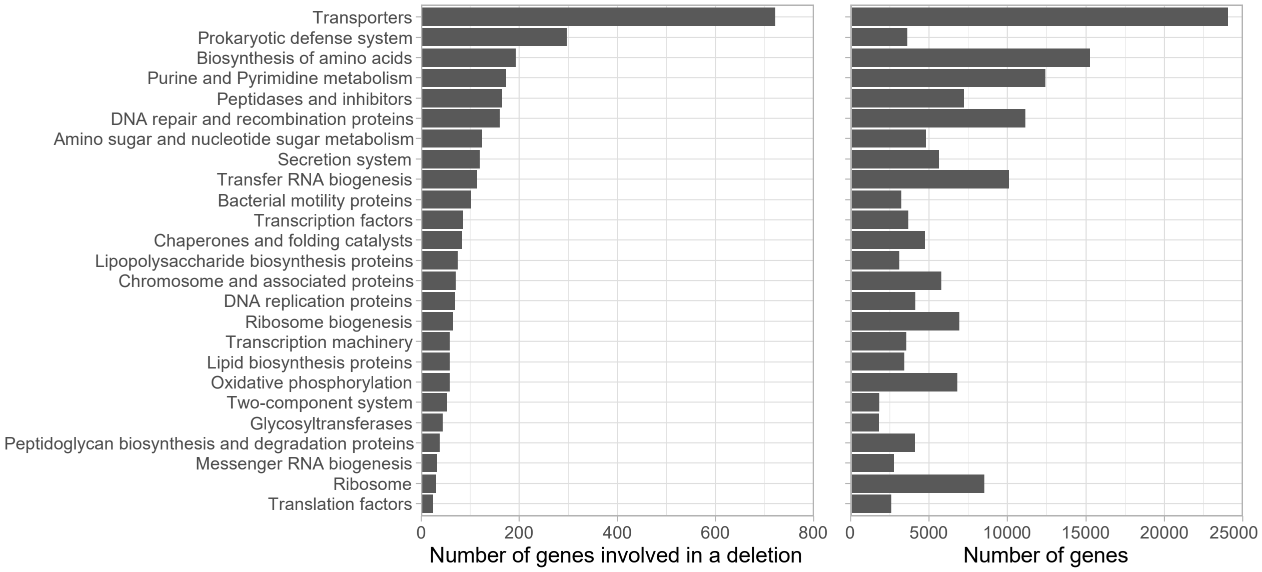


**Figure S5.** The number of genes in total (right) and those involved in a deletion (left) in each gene functional category among the 178 rMAGs analyzed. Gene categories are sorted by the number of genes involved in deletions.

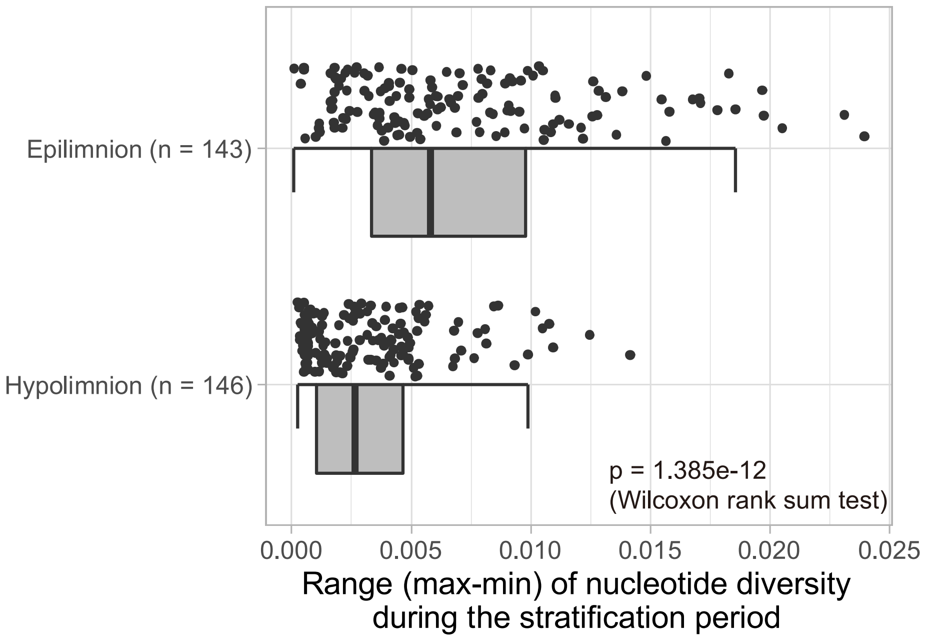


**Figure S6.** Distribution of the range (maximum value–minimum value) of the nucleotide diversity of each rMAG (represented by each point) during the stratification period (May to December). The rMAGs for which the nucleotide diversity could be calculated for more than four out of the eight months in each of the water layers were included in the analysis. The range were significantly broader in the epilimnion than in the hypolimnion, according to the Wilcoxon rank sum test.

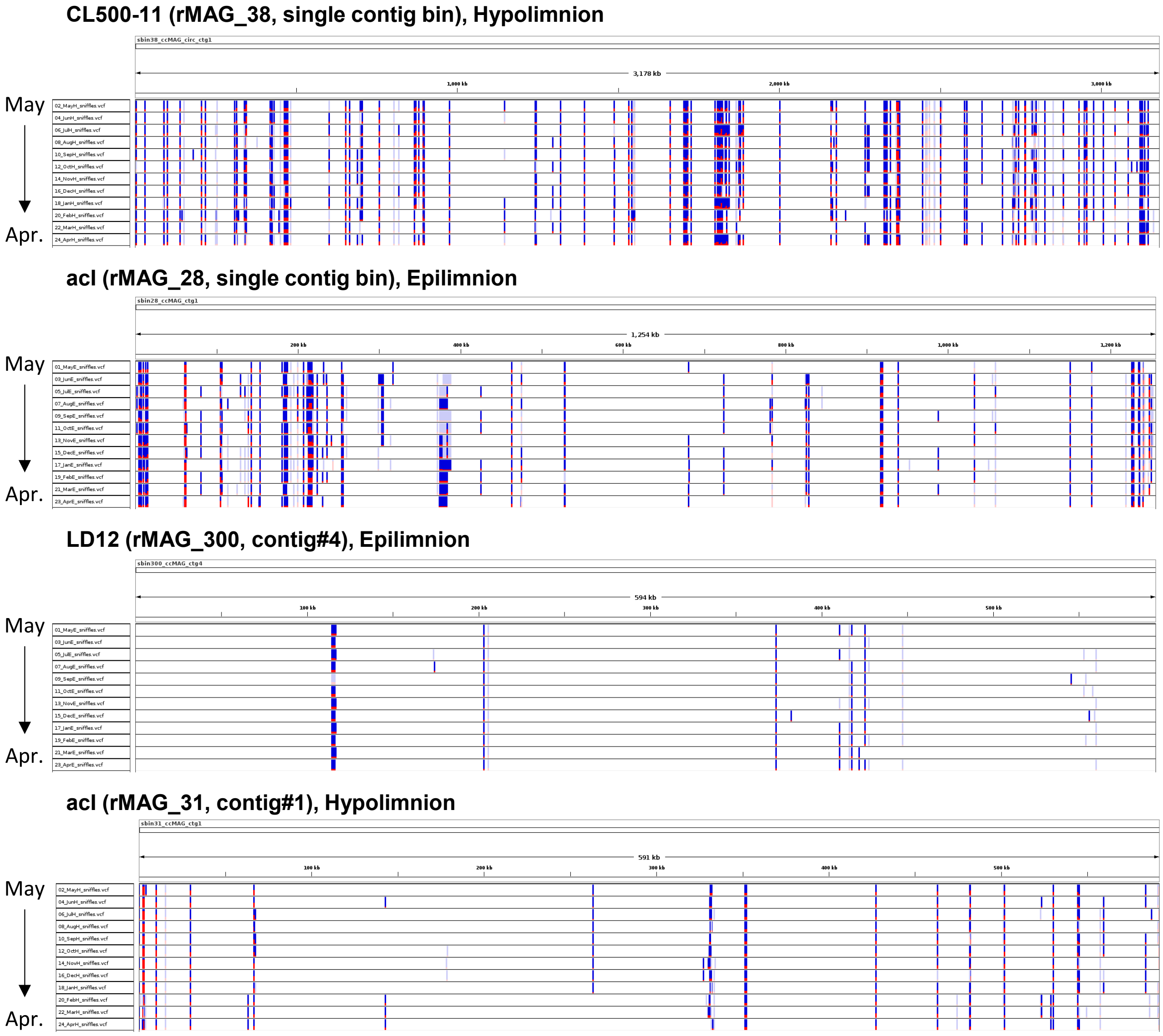


**Figure S7.** Temporal profile of SVs in four representative rMAGs that showed continuously high read coverage during the study period in either of the water layers. The top two rMAGs are single-contig bins while the longest contig was displayed for the other two rMAGs. SV positions in each sequence were visualized with Integrative Genomics Viewer (IGV) and chronologically sorted from top (May 2018) to bottom (April 2019). The allele frequency of each SV was represented as the height of the red-colored part against the blue-colored part. Shaded SVs are those filtered out by Sniffle software due to poor or inconsistent support with read mapping results. Most SVs were continuously present during the study period.

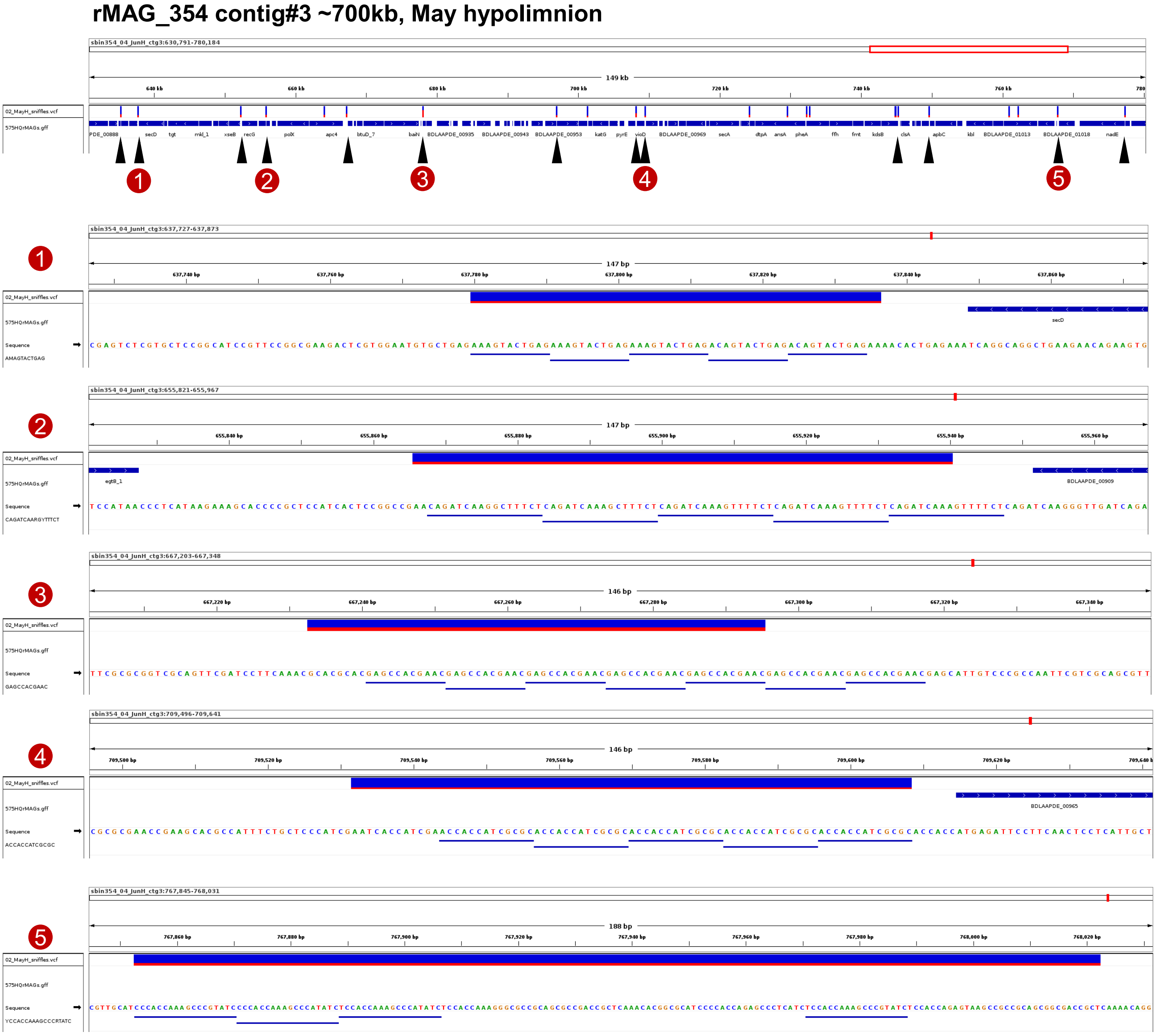


**Figure S8.** Deletions on intergenic tandem repeats in rMAG_354 (Phycisphaerales of the phylum Planctomycetota). The top panel indicates the SV profile (visualized by IGV in the same manner as Fig. S7) in a representative genomic region of the rMAG. The second row in the panel shows ORFs. Black arrows indicate intergenic deletions; red circled numbers indicate those involve tandem repeats, for which enlarged visualizations are shown in the bottom panels. Nucleotide sequences and repeat motifs are shown in the third and fourth rows in the enlarged panels.

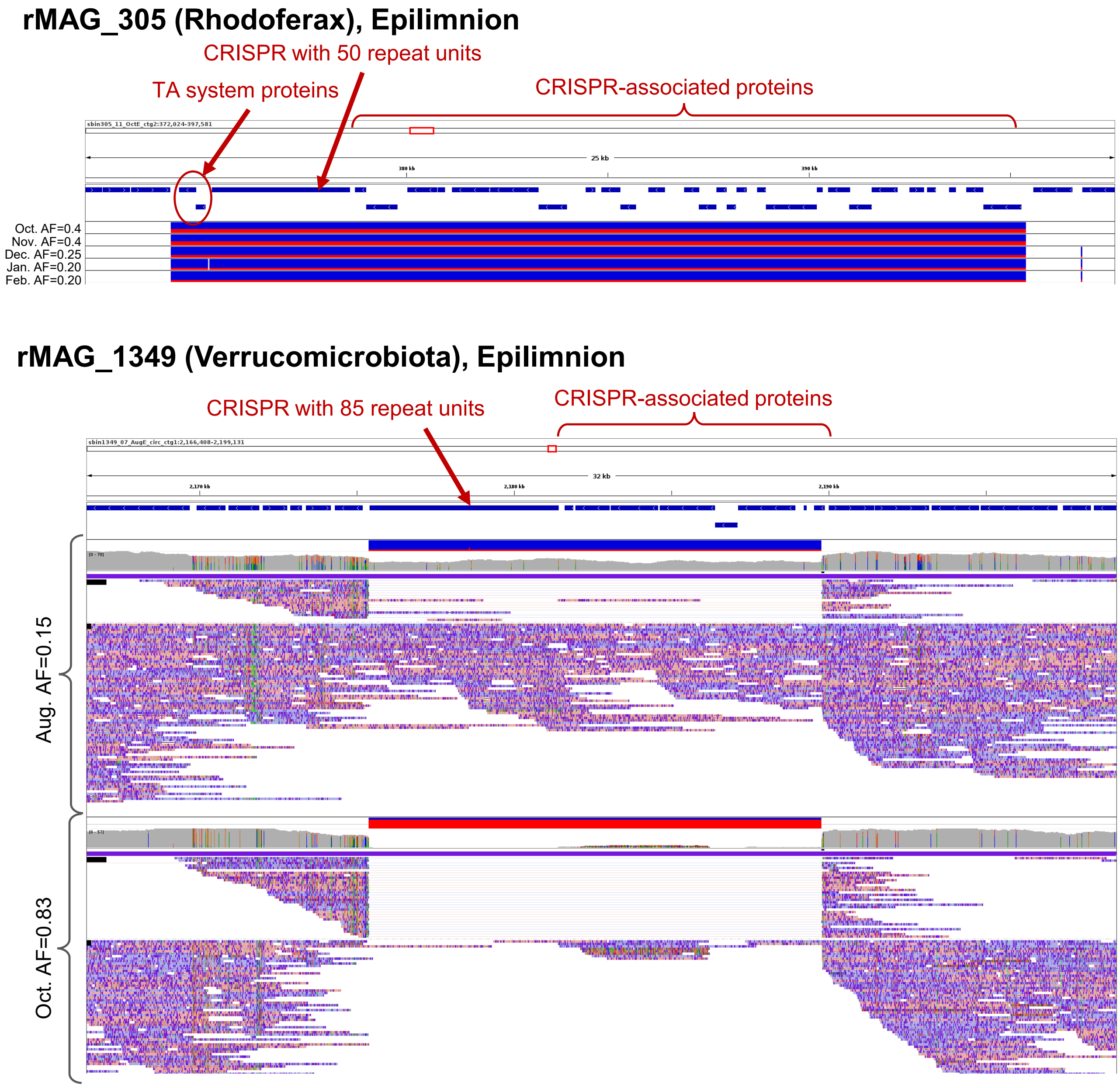


**Figure S9.** Deletions involving a CRISPR–Cas system. ORFs and SVs (all are deletions here) are visualized by IGV in the same manner as Figures S7 and S8. In the top case (rMAG_305), the deletion was continuously detected from October to February in the epilimnion, with decreasing allele frequency. The deletion also included TA system proteins downstream of the CRISPR. In the bottom case (rMAG_1349), the allele frequency shifted more quickly from 0.15 in August to 0.83 in October. Pileup and coverage tracks of mapped long reads were shown for each sample and indicated that many reads were aligned to the SV region in August while most of the reads bridged the edges of the SV region in October. AF, allele frequency.

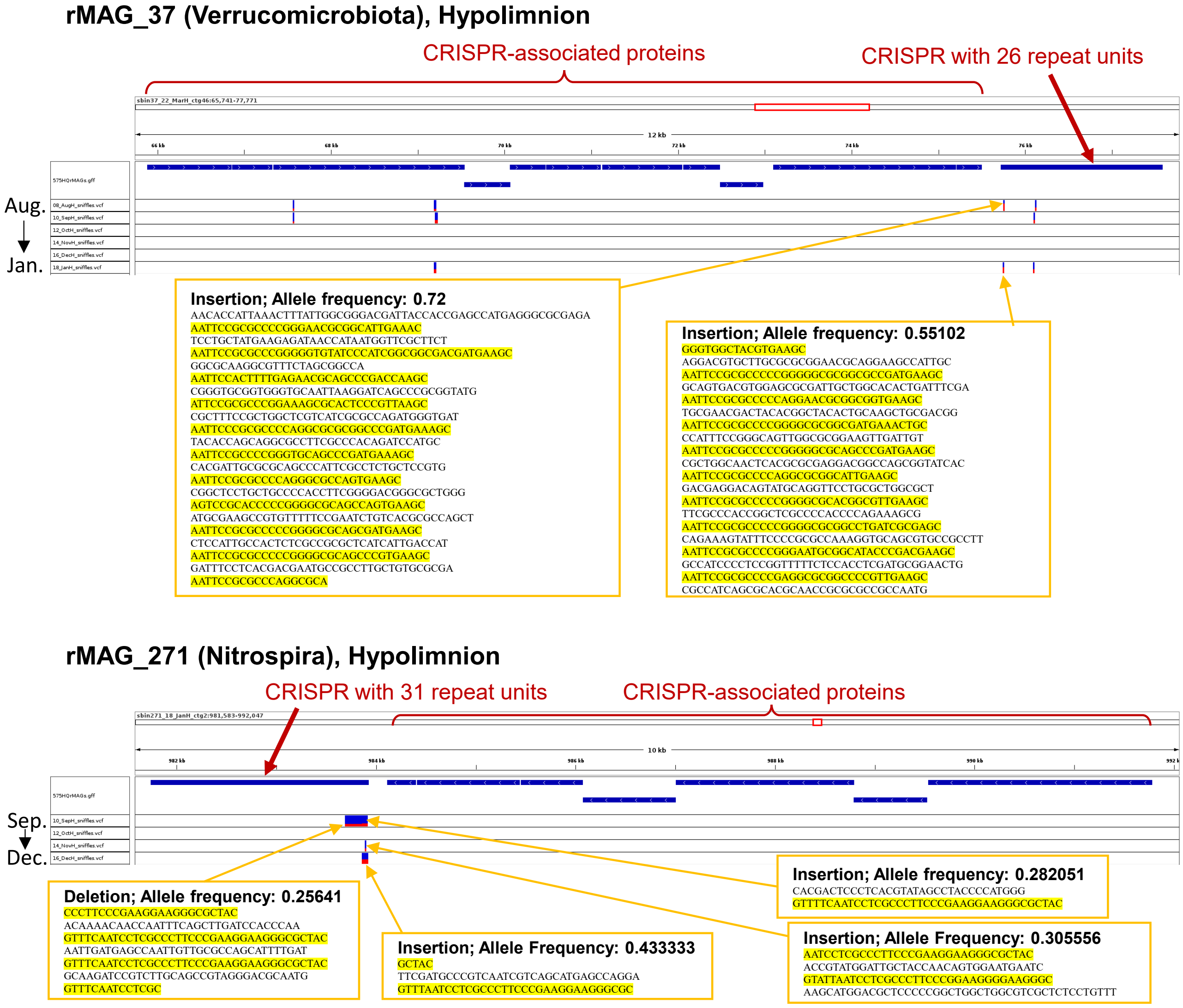


**Figure S10.** SVs associated with variation in CRISPR spacer sequences. ORFs and SVs are visualized by IGV in the same manner as Figures S7 and S8. Both rMAGs showed a shift of SV pattern in the CRISPR sequences during consecutive months in the hypolimnion. SV type, allele frequency, and the sequence involved in each SV are shown in an orange box, in which CRISPR repeat sequences are shaded yellow. Note that sequences shown for insertion were predicted using sequences of the mapped raw long reads and thus unpolished (i.e., error-prone).

### Captions for other Supplementary Materials

**Supplementary Table S1**

Environmental parameters and sequencing statistics of the 24 samples.

**Supplementary Table S2**

Statistics and analytical results for each of the 575 rMAGs. Note that those with > 10× short-read coverage in the representative sample (n = 178) were mainly analyzed for their microdiversity (see column S in the Excel sheet).
